# Oligodendrocyte progenitor proliferation is disinhibited following traumatic brain injury in LIF heterozygous mice

**DOI:** 10.1101/2020.12.09.418343

**Authors:** Michelle J. Frondelli, Marie. L. Mather, Steven W. Levison

## Abstract

Traumatic brain injury (TBI) is a significant problem that affects ∼500,000 children each year. As cell proliferation is disturbed by injury and is required for normal brain development, we investigated how a pediatric closed head injury (CHI) would affect the progenitors of the subventricular zone (SVZ). Additionally, we evaluated the contribution of Leukemia Inhibitory Factor (LIF) using LIF-heterozygous mice (LIF Het), as LIF is an injury-induced cytokine, known to influence neurogenesis and gliogenesis. CHI’s were performed on P20 LIF Het and WT mice. Ki-67 staining and stereology revealed that cell proliferation increased ∼250% in injured LIF Het mice compared to the 30% increase observed in injured WT mice at 48 h post CHI. Furthermore, Olig2+ cell proliferation increased in the SVZ and white matter of LIF Het injured mice at 48 h recovery. Using an 8-color flow cytometry panel, the proliferation of three distinct multipotential progenitors were greater in LIF Het injured mice compared to WT injured mice. Early oligodendrocyte progenitor cell (OPC) proliferation was 6-fold higher in LIF Het injured mice compared to WT injured mice. In vitro, addition of LIF decreased overall cell proliferation and OPC proliferation compared to controls. Addition of LIF to OPC cultures induced an increase of phospho-Akt after 20 minutes and an increase of phospho-S6RP at 20 and 40 minutes of exposure, suggesting that LIF stimulates the mammalian target of Rapamycin pathway. Altogether, our data provide new insights into the regulatory role of LIF in suppressing neural progenitor cell proliferation after a mild TBI.

**Main Points:**

- OPC proliferation is dis-inhibited in LIF haplodeficient mice.
- LIF directly inhibits glial progenitor cell proliferation.
- LIF stimulates the mTOR pathway.

## Introduction

Traumatic brain injury (TBI) affects 2.87 million people annually in the United States alone and of these, 30% occur in children under the age of 14. Of the roughly 860,000 children who sustain a TBI, ∼10% are hospitalized, 2% die and 90% require an immediate emergency room visit. Including the direct and indirect medical costs, TBI accounts for a lifetime economic cost of $ 76.5 billion dollars (Centers for Disease Control and Prevention (CDC), 2019). TBI in children is especially consequential because it can change the normal, proliferative process which in turn, will alter brain development, impair cognition, create learning difficulties, affect motor skills and disturb emotional behavior. Therefore, there is an essential need to establish how neurotrauma affects normal brain development and what can be done to mitigate the loss of newly created cells.

A number of cytokines are produced subsequent to a TBI that participate in tissue remodeling. Among them is Leukemia Inhibitory Factor (LIF), which is a cytokine that binds with high affinity to a heterodimeric receptor consisting of the LIF receptor and glycoprotein 130. LIF levels surge as a result of brain injury with increased levels reported after cerebral ischemia, stroke, multiple sclerosis, Alzheimer’s disease, Parkinson’s disease, controlled cortical impact (CCI) and seizure (Buono et al., 2015; Deverman & Patterson, 2012; Felling et al., 2016; Goodus et al., 2016; Soilu-Hanninen et al., 2010; Vanderlocht et al., 2006). LIF reduces injury progression through its neurotrophic actions, promotes remyelination, and activates astrocytes (Deverman & Patterson, 2012; Felling et al., 2016; Goodus et al., 2016; Gresle et al., 2012). Additional studies have found that LIF reduces neurogenesis in the olfactory bulb and subventricular zone (SVZ) by acting directly on neural stem cells (NSCs), thus promoting NSC self-renewal, leading to an expansion of the NSC pool (Bauer & Patterson, 2006; Buono et al., 2012). These studies establish LIF as an important mediator of injury repair; however, none of these studies address the role of LIF in regeneration from TBI especially in the context of pediatric TBI, where the potential for regeneration is high (Covey et al., 2010).

The SVZ harbors NSCs and a variety of progenitors and is quite active in the pediatric brain. As the pediatric brain is still growing, the SVZ is very active and continuously making new cells. Injuries stimulate cell proliferation within the SVZ, as shown using a variety of pediatric brain injury models including CCI (Goodus et al., 2015; Ramaswamy et al., 2005); hypoxia-ischemia (Buono et al., 2015; Ong et al., 2005; Plane et al., 2004) and penetrating lesions (Covey et al., 2010; Jinnou et al., 2018).These newly produced cells have the regenerative potential to differentiate into neurons and glia (Faiz et al., 2005; Kernie & Parent, 2010; Lois & Alvarez-Buylla, 1993; Parent et al., 2002). Although studies have shown a strong correlation between SVZ proliferation and injury induced LIF, they have also shown model-dependent differences in the SVZ proliferative response. Moreover, despite the high incidence of closed head injuries (CHIs), few studies have evaluated the effects of CHIs on the dynamics of cell proliferation within the SVZ. In a previous study, we found that LIF Heterozygous (LIF Het) mice experienced more axonal damage than wild-type (WT) mice with persisting behavioral deficits. Therefore, in the studies described here, we hypothesized that a mild CHI would alter the composition and proliferation of the NSCs and progenitors of the SVZ and that these responses would be diminished in mice deficient in LIF, which in turn would contribute to the failure of recovery.

## Materials and Methods

### Closed Head Injury (CHI)

The animal protocol for the work described in this report was approved by the New Jersey Medical School IACUC (#999900841) and these studies were in accordance with the National Institute of Health Guide for the Care and Use of Laboratory Animals (NIH Publications No. 80-23) revised in 1996. We further attest that all efforts were made to minimize the number of animals used and that efforts were made to ensure minimal suffering. To simulate a pediatric brain injury, mice were injured by CHI on CD-1, P20 mice, the age roughly equivalent to humans aged 2-3 years old (Semple et al., 2013). LIF levels reach a peak at 48 h post injury (PI), therefore, both WT and LIF Het mice were injured by CHI and terminated 48 h later (Goodus et al., 2016). LIF heterozygous mice were produced from a mouse line in which gene encoding LIF was mutated by disrupting the 3rd exon that truncated the LIF protein so that it was missing the last carboxyterminal 81 amino acids, including the last nine, that are essential for its biological activity (Stewart et al., 1992). The eCCI Model 6.3 piston driven impactor (Custom Design & Fabrication, Cat. #23298-047) was used to produce a bilateral CHI midway between Bregma and Lambda. Mice were anesthetized and an incision along the midline was created to expose the skull. A round, metal impactor tip of 5 mm in diameter was used to create the CHI directly on the intact skull. The impactor was driven onto the skull at a velocity of 4m/s to a depth of 1mm past the zero point on the skull surface with a dwell time of 150 ms. Age-matched sham-operated animals received the same isoflurane treatment and scalp incisions. Mice were monitored continuously for 2 h after surgery and received buprenorphine subcutaneously as post-operative care at 0.05 mg/kg. Mice were returned to their cages and checked daily until terminated.

### Intracardiac Perfusions

Two days following CHI, mice were anesthetized with a mixture of ketamine (75 mg/kg) (Henry Schein, Cat. #11695-0702) and xylazine (5 mg/kg) (Akorn Animal Health, Cat. #59399-111-50) and then perfused with RPMI media containing 5 U/mL heparin (Sagent Pharmaceuticals, Cat. #NDC 25021-400-10) followed by 3% paraformaldehyde (PFA) (Electron Microscopy Science, Cat. # 15714-S) in PBS. Brains were dissected; post-fixed overnight in 3% PFA/PBS, and cryoprotected for 24 h in 30% sucrose in water. After 24 h, the brains received fresh sucrose for 4 h and then the brains were frozen in Tissue-Tek OCT matrix embedding medium (Sakura Finetek, Cat. #4583).

### Coronal Sections and Immunostaining

The brains were sectioned coronally at 40μm containing the SVZ using a Leica CM 1950 cryostat (Leica, Cat. #CM1510 S). Sections were washed twice with TBS for 5 m and incubated for 30 m with 0.3% Triton-X (Fisher Scientific, Cat. #9002-93-1) in tris-buffered saline (TBS). Sections were washed twice with TBS for 5 m and blocked with 10% goat superblock for 1 h followed by incubation with primary antibodies in 1% goat serum/0.05% Triton X-100/TBS at 4°C overnight. Sections were washed three times with 0.05% Triton X-100/TBS for 30 m followed by one 5 m wash in TBS and incubated with secondary antibodies with 1% goat serum for 2 h at 37°C. Sections were washed three times with 0.05% Triton X-100/TBS followed by one wash in TBS for 5 m. Coverslips were affixed using Prolong Gold Antifade Mountant with 4’,6-diamidino-2-phenylindole (DAPI) (ThermoFisher Scientific, Cat. # P36935).

The following primary antibodies were used: rabbit anti-Ki-67 (1:50; Abcam, Cat. #ab15580, RRID: AB_443209); rat anti Ki-67 (1:250, eBioscience, Cat. #14-5698-82, RRID: AB_10854564); rabbit anti-Olig2 (1:250, Millipore Sigma, Cat. # AB9610, RRID: AB_570666). The following secondary antibodies were used: Alexa 488 conjugated donkey anti rabbit (Cat. #711-485-152, RRID: AB_2492289); Cy3-conjugated donkey anti rat (Cat. #712-165-153, RRID: AB_2340667); all used at 1:200 dilution and purchased from Jackson ImmunoResearch.

### Stereology

To quantify double-immunofluorescently labeled cells, sections were analyzed using the MicroBrightField Stereo Investigator program on an Olympus BX51 microscope. Under the MicroBrightField Optical Fractionator workflow, each SVZ and white matter (WM) region was traced and randomly placed counting frames were applied to sample colocalized markers within the traced region at predetermined regular X and Y intervals within volume samples. Twenty percent of the SVZ and WM region were counted and 50% of the subgranular zone (SGZ) was counted, as determined necessary by using cavalieri point counting equations including profile area and profile boundary equations. Image stacks were captured at 100X using an immersion oil objective. Selection criteria for counting an object within the sampling frame were implemented using stereological counting rules. Inclusion and exclusion counting criteria were followed and recorded only when cells were double positive for Ki-67 and Olig2.

### Flow Cytometry (In Vivo)

After CHI, mice were provided drinking water containing EdU that they could drink ad libitum for 48 h. The water contained 1 mg/mL 5-ethynyl-2-deoxyuridine (EdU) (Thermo Fisher Scientific, Cat. # E10187) and 1% sucrose. After two days, mice were terminated and both hemispheres of the SVZ were isolated by microdissection, pooled and incubated with 26 Wünsch units/mL of Liberase DH (Millipore Sigma, Cat. #5401054001) and 40 µg/ml DNase 1 (Millipore Sigma, Cat. # D5025-150KU) in PBS with 2.8 mM MgSO_4_ and shaken at 24 RPM at 37°C for 45 m. Enzymatic digestions were quenched with 8 mL of PGB and 0.006% HEPEs (Corning, Cat. #25-060-Cl) and the cells were centrifuged for 5 m at 300 x g. The cells were dissociated by repeated trituration, collected and counted. Cells were diluted to 1 million cells/mL in PGB.

For surface marker analysis, the cells were incubated in PGB for 25 m with fluorescent-conjugated antibodies against: CD15-v450, CD133-SB600, CD140a-PE, NG2 chondroitin-A700 sulfate proteoglycan, GLAST, and O4-APC (Table 1). The cells were washed with PGB at 300 g and incubated with secondary antibodies and Live/Dead blue in PGB for 25m and then washed three times in PBS/ dextrose/BSA (PGB) at 300 g. The cells were fixed with 1% PFA for 20 m, collected, resuspended and stored at 4°C overnight.

**Table 1:** List of primary and secondary antibodies used to complete flow cytometry studies.

| Antibody/Reagent | Dilution | Host | Isotype | Fluorophore | Clone | Catalog # | Vender | RRID |
| --- | --- | --- | --- | --- | --- | --- | --- | --- |
| LIVE/DEAD | 1:1000 | NA | NA | Blue | NA | L34961 | Thermo Fisher | NA |
| CD15 (LeX) | 1:20 | Mouse | IgM | V450 | MC480 | 561561 | BD Biosciences | AB_10926202 |
| CD140 (PDGFRα) | 1:160 | Rat | IgG2a, K | PE | APA5 | 135906 | Biolegend | AB_1953269 |
| NG2 | 1:50 | Rabbit | IgG | NA | NA | AB5320 | Millipore | AB_91789 |
| Goat anti Rabbit | 1:100 | Goat | NA | A700 | NA | A21038 | Life Technologies | AB_1500674 |
| GLAST (biotin) | 1:11 | Mouse | IgG2a, K | NA | ASCA-1 | 130-118-984 | MACs (Miltenyi) | AB_2733473 |
| Streptavidin APC | 0.125 ug/ test | NA | IgG2a, K | 780 | NA | 47-4317-82 | eBioscience | AB_10366688 |
| O4 | 1:125 | Mouse | IgG1 | APC | REA576 | 130-119-897 | MACs (Miltenyi) | AB_2751913 |
| CD133 | 0.01 ug/ test | Rat | IgG1, K | Superbright 600 | 13A4 | 63-1331-80 | Thermo Fisher | AB_2734873 |
| FcR Block | 1:20 | NA | NA | NA | NA | 30-092-575 | MACs (Miltenyi) | NA |
| Click-it Plus EdU<br>Alexa Fluor 488 Flow<br>Cytometry Assay Kit | NA | NA | NA | 488 | NA | C10633 | Thermo Fisher | NA |

The following day, the cells were collected by centrifugation at 300 g, and manufacturer instructions were followed to label and detect EdU+ cells. All sample data were collected on a BD Fortessa flow cytometer (BD Biosciences Immunocytometry Systems). Matching isotype controls for all antibodies were used and gates were set based on these isotype controls. Data were analyzed using FlowJo software (Tree Star, Inc.). Live, dead, and EdU-controls were used to determine gating strategy. Gates were set according to a 98% confidence interval on the isotype controls.

### Neurosphere Cultures

Neonatal WT, CD1 mouse pups at P4-P5 were decapitated and their brains were immediately removed. The SVZ was dissected out of each hemisphere and the tissue was digested with a solution containing 0.1% trypsin (Invitrogen, Cat. #15400054), 10 units/mL papain (Worthington Biochemical Corporation, Cat. # LS003124), 40 μg/mL DNase I and 3 mM MgSO_4_ in DMEM/F-12 for 5 m at 37°C with intermittent shaking. The enzymatic reaction was stopped using an equivalent volume of 10% newborn calf serum in DMEM/F-12. The tissue was centrifuged at 100 x g for 5 m followed by a wash with DMEM/F-12, after which the pellet was resuspended in 3 mL of DMEM/F-12, mechanically triturated until smooth and then filtered through a 40 μm cell sieve. After centrifugation, the cells were plated at a density of 5 x 10^5^ cells/mL of neurosphere growth medium with 200 μl/cm^2^ in 60 mm tissue culture dishes. The neurosphere growth medium consisted of DMEM/F-12 supplemented with 15 mM HEPES, 1% GlutaGro (Corning, Cat. #25-015-Cl), 1% Antibiotic-Antimycotic (Thermo Fisher, Cat. #1540062), 2% B27 supplement (Thermo Fisher, Cat. #17504001), 20 ng/mL recombinant human epidermal growth factor (EGF) (PeproTech, Cat. # AF-100-15) and 10 ng/mL recombinant human fibroblast growth factor-2 (FGF-2) (PeproTech, Cat. #100-18C) and 1 ng/mL heparin sulfate.

Cells were propagated at 37°C with humidified 5% CO_2_ to permit primary neurosphere formation and were fed every two days. Primary neurospheres were collected after seven days and were dissociated for 5 m at 37°C in 70% Accutase (Millipore Sigma, Cat. # SCR005) in DMEM/F-12 and plated onto 60 mm tissue culture dishes as described above. Cells were fed every two days until they reached ∼200 μm in diameter after roughly 5 days in vitro (DIV) after passaging. For cell cycle studies, the cells were deprived of growth factors for 15 h. Growth factors (EGF & FGF-2) were reintroduced to the cultures ± recombinant murine LIF (PeproTech, Cat. #250-02) at 5 ng/mL. After 24 h, EdU was added to cell cultures for 24 h at 10 μM with the exception of one plate that was deprived of EdU, thus serving as a negative control.

### Flow Cytometry (In Vitro)

Cells were collected and incubated with 2.6 Wünsch units/mL of Liberase DH and 20μg/mL DNAse in DMEM/F-12 and incubated at 37°C for 5 m with intermittent stirring. Enzymatic digestions were quenched with equal volume of DMEM/F-12 containing 20μg/mL DNAse. The cells were centrifuged for 5 m at 300 g and were dissociated by repeated trituration. Cells were collected by centrifugation, counted, diluted to 1 million cells/ mL in of DMEM/F-12 and stained according to previously described methods (See “Flow Cytometry (*In Vivo*).)

### OPC Cultures

Newborn WT, CD1 mouse pups at P1 were decapitated and their brains were immediately removed. The skulls were cut along the longitudinal fissure and the olfactory bulbs, cerebellum and meninges were removed from each brain. The remaining tissue was enzymatically digested with a solution containing 0.25% trypsin (Invitrogen, Cat. #15400054), and 40 μg/mL DNase I in PBS with 2.8 mM MgSO_4_ for 15 m at 37°C with intermittent shaking. The enzymatic reaction was stopped using an equivalent volume of a solution containing 10% fetal bovine serum, 1% Glutagro (Corning, Cat. #25-015-Cl), 1% Pen Strep (Corning, Cat. #30-002-Cl), 0.6% glucose in Eagle’s Minimum Essential Medium (Gibco, Cat. #11090-081) (MEM-C). The tissue was triturated and passed through a 100 μm cell sieve. The tissue was shaken at 1500 rotations per minute (RPM) for 8 m followed by resuspension in MEM-C and filtered through a 40 μm cell sieve. Cells were counted and plated at a density of 1.5 x 10^7^ cells per 75 cm^2^ flask in MEM-C. Media was replaced after 4 DIV and then replaced every two days after being maintained at 20% O_2_, 5% CO_2_, & 37°C.

At 9 DIV, OPCs were amplified by shaking the cells at 200 RPM for 1 h at 37°C to remove microglia. Media was removed and replaced and OPCs were placed into an cell culture incubator for 1 h at 37°C. OPCs were shaken for 18 h at 260 RPM at 37°C. Supernatant was passed through a 20 μm Nitex screen and centrifuged at 150 x g for 5 m. Supernatant was removed and pellet was resuspended in 5 mL of solution containing 2% B27, 1% Pen Strep, 1% GlutaGro & 0.5% fetal bovine serum in DMEM/F12-HEPEs (N2B2). Cell suspension was plated onto a 100 mm bacteriological plastic dish for 15 m at 37°C to allow microglia to adhere to the dish. Cell suspension was collected, dishes were rinsed with N2B2, and cells were centrifuged at 150 x g for 5 m & resuspended in 33% B104 conditioned N2B2 & 67% N2B2 supplemented with 100ng/uL FGF-2 (N2S). Cells were counted and plated onto fibronectin-coated wells at a density of 3 x 10^4^ cells/cm^2^ in N2S. Cells were maintained in a three gas incubator with 2%O_2_, 5% CO_2_ and 93% N_2_ at 37°C. Cells were fed every two days by replacing 2/3 of the media.

### Western Blots

OPC cultures were washed once with PBS. Lysis buffer containing protease and phosphatase inhibitors (Thermo Fisher, Cat. #89900) was applied to cell cultures for 10 m on ice. OPCs were scraped, collected, and incubated on ice for an additional 10 m. Samples were sonicated 3 times using 5s pulses and centrifuged at 9300 x g for 5 m. The supernatant was transferred to a fresh tube and protein was quantified using a Pierce BCA Protein assay kit (Thermo Fisher; Cat. #23225).

For Western blot analyses, 30 μg of denatured protein was loaded onto a 4-12% Bis-Tris polyacrylamide gel (Invitrogen, Cat. #NP0321PK2). Five μL of Amersham ECL Rainbow Marker was included as a molecular weight standard (Millipore Sigma, Cat. #GERPN800E). The proteins were separated by electrophoresis at 200V, 130mA and 90W for 1 h. Proteins were transferred onto nitrocellulose at 30V, 180mA and 8W for 1 h and 5 m. The membrane was incubated for 1 h with 5% bovine serum albumin in 1 x TBS/0.1% Tween (TBST) before being incubated with primary antibodies: pAkt (1:2000, Cell Signaling Technology, Cat. #4060, RRID: AB_2315049), and pS6RP (1:2000, Cell Signaling Technology, Cat. #4858S, RRID: AB_916156). Membranes were probed for primary antibodies overnight at 4°C, washed with 1 x TBST, and incubated with Anti-rabbit IgG, HRP-linked Antibody (1:2500, Cell Signaling Technology, Cat. #7074P2, RRID: AB_2099233) for 90 m with agitation, washed, and bands visualized using Western Lightning chemiluminescence reagent (PerkinElmer, Wellesley, MA, Cat. #NEL103E001EA). Imaging was performed using a BioRad ChemiDoc Imaging System combined with Image Lab software. Membranes were washed three times in TBST and stripped of antibodies using a buffer containing 0.625 M Tris-HCL, 10% sodium dodecyl sulfate and 0.7% 2β- mercaptoethanol in water for 30 m with agitation at 50°C. Membranes were rinsed once for 5 m with TBST and incubated for 1 m in chemiluminescence reagent to ensure that original signal was removed. The membrane was washed again 4 times in TBST for 5 m each and blocked again before incubating with primary antibodies: total Akt (1:1000, Cell Signaling Technology, Cat. #9272S, RRID:AB_329827) & total S6RP (1:1000, Cell Signaling Technology, 2217, RRID:AB_331355) overnight at 4°C. Washes, secondary incubation, chemiluminescence incubation and imaging steps were repeated.

### Statistical Analyses

Statistical analyses were performed using GraphPad Prism software. The means between two groups were compared using two-tailed, unpaired t test. The means between more than two groups were compared using one-way ANOVA with Tukey’s *post hoc* test. Error bars denote standard deviations (SD). Comparisons were interpreted as significant when associated with p < 0.05.

## Results

### Cell proliferation is more robust in LIF Het mice after CHI

The SVZ and SGZ demonstrate enhanced neurogenesis and excitability after moderate to severe traumatic brain injuries. Since LIF is a key niche factor within these germinal zones, we also examined the effects of pediatric CHI on the proliferative cells of the SVZ *in vivo* in both WT and LIF Het mice. To evaluate changes in neurogenesis and gliogenesis in these regions pediatric aged mice (P20) were administered a CHI halfway between Bregma and Lambda and terminated by intracardiac perfusion two days later (Fig. 1A). Cryosections were collected to span the majority of the SGZ and the SVZ and they were immunolabelled for Ki-67, as a marker of proliferation. Stereological counts of Ki-67+ cells in the SGZ from WT sham, WT injured and LIF Het injured mice did not show statistical differences between the groups (Supplemental Figure 1). Therefore, we focused our efforts on defining changes within the SVZ.

**Figure 1.**
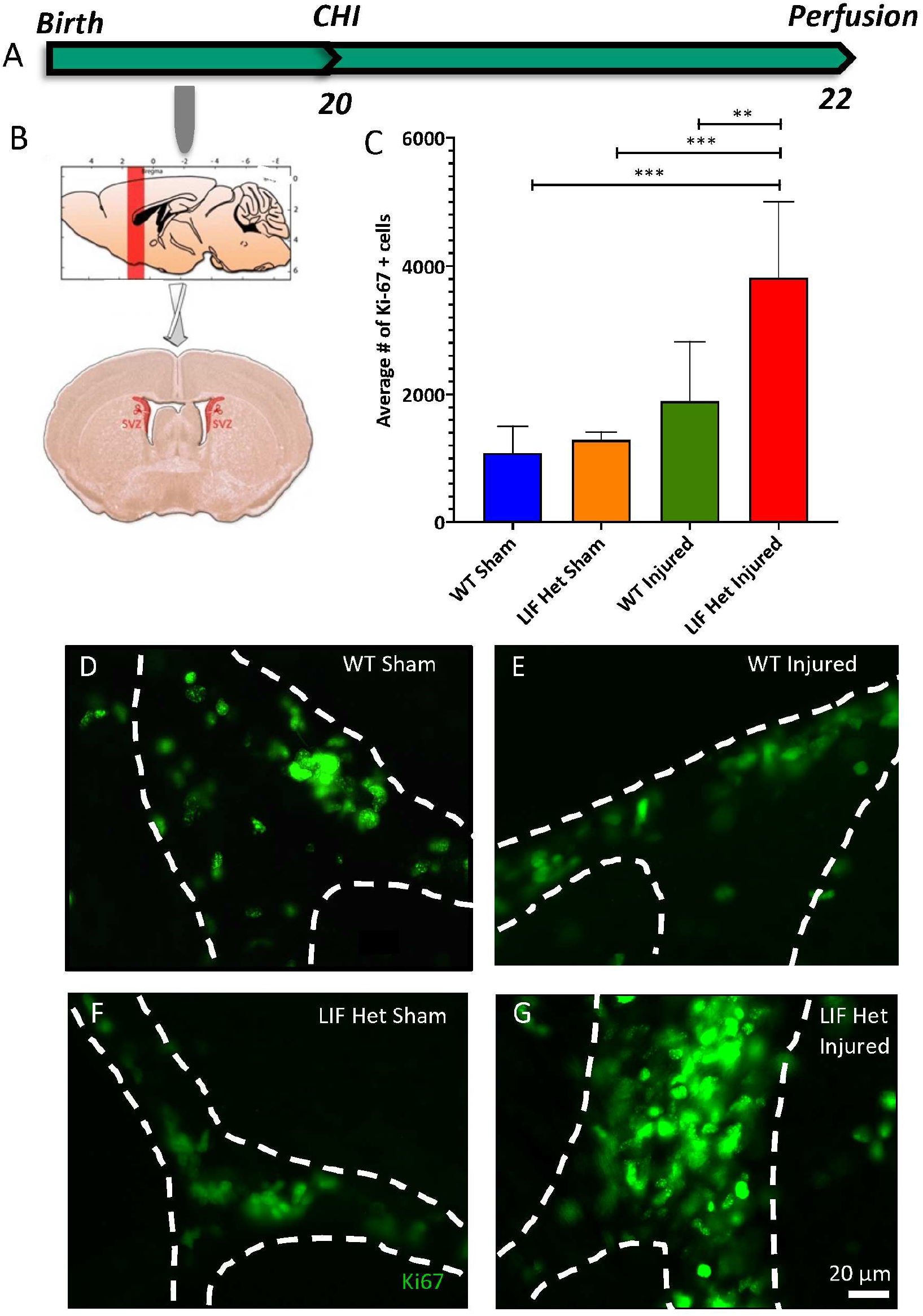
SVZ cell proliferation is more robust in LIF Het mice after a CHI. (A) Schematic of experimental design. WT and LIF Het mice were injured at postnatal Day 20 and analyzed 2 DPI. (B) Images depicting the level of the brain below the impactor in sagittal and coronal sections. (C) Coronal 40 μm sections containing the SVZ were stained for Ki-67. Stereological counts of Ki-67+ cells were obtained in the SVZ from LIF Het CHI, LIF Het sham, WT sham, and WT CHI mice. Values represent the means ± SDs from n=5 animals per group. **p≤0.01 vs. WT injured; ***p≤0.001 vs. LIF Het injured, LIF Het sham by one-way ANOVA and Tukey’s *post hoc* test. (D-G) Representative images of the dorsomediolateral SVZ below the center of the impact zone 2 DPI from (D) WT sham, (E) WT injured, (F), LIF Het sham, and (G) LIF Het injured mice. Dotted lines outline the SVZ. Magnification bar = 20 µm.

Stereological cell counts of Ki-67+ cells were performed on both hemispheric SVZ’s of each of three sections (0.48 mm apart) and averaged. LIF Het injured mice revealed a 3-fold increase in Ki-67+ cells over LIF Het sham and WT sham mice (p≤0.001) and a 2-fold increase in the average number of Ki-67+ cells when compared to the WT injured mice (p≤0.01) at 2 days post injury (DPI) (Fig. 1C). Figure 1D-G depict representative images that correlate with the stereological quantification of Ki-67+ cells for WT sham (Fig. 1D), WT injured (Fig. 1E), LIF Het sham (Fig. 1F) and LIF Het injured mice (Fig. 1G). Altogether, these data indicate that LIF deficiency enhances the normal, proliferative response to pediatric CHI.

### Oligodendrocyte progenitor proliferation is more robust in LIF Het mice after CHI

As there were more proliferating cells in the LIF Het injured mice than WT injured and sham controls at 48 h post CHI (Fig. 1C), and as there was no difference in the numbers of proliferating neuroblasts (Supplemental Figure 2), we next evaluated the number of Ki-67+/Olig2+ cells in the SVZ at 48 h post CHI. While a previous study had shown that at early time points of recovery from a controlled cortical impact injury in mice that the majority of proliferating cells were NG2 (Susarla et al., 2014) we chose to stain the cells using Olig2, as this transcription factor is expressed throughout the entire oligodendrocyte lineage including the progenitors, pre-oligodendrocyte, immature oligodendrocyte and mature oligodendrocyte stages (Rowitch et al., 2002). Figure 2A-D depict representative images that correlate with stereological quantification of Ki-67+/Olig2+ cells for WT sham (Fig. 2A), WT injured (Fig. 2B), LIF Het sham (Fig. 2C) and LIF Het injured mice (Fig. 2D). At 2 DPI, LIF Het injured SVZs had a 3-fold increase of Ki-67+/Olig2+ cells than the WT injured mice, LIF Het sham mice, and WT sham mice (p≤0.0001) (Fig. 2E).

**Figure 2.**
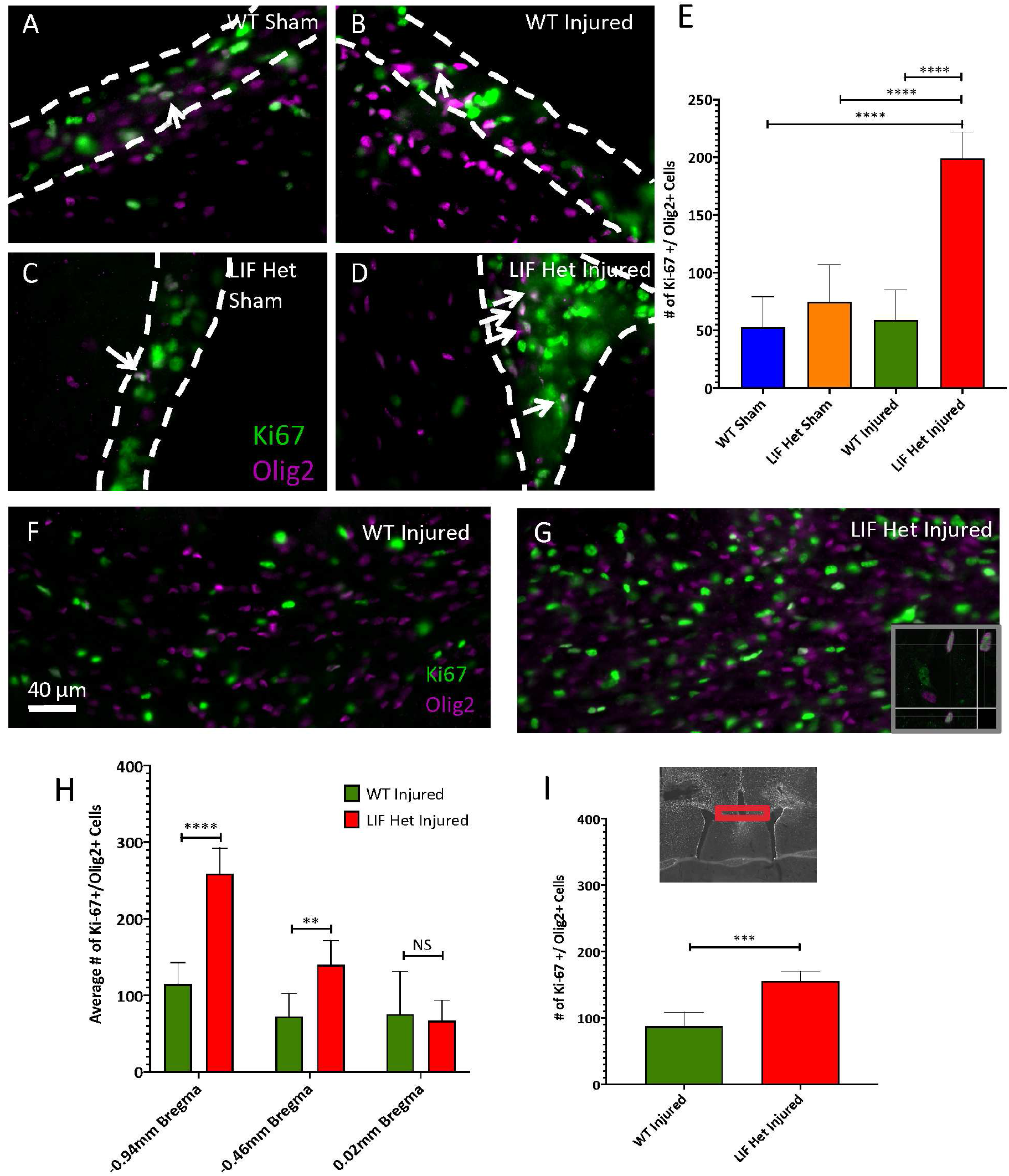
Oligodendrocyte progenitor proliferation is more robust in LIF Het mice after CHI. (A-D) Representative images of the SVZ 2 DPI from (A) WT sham, (B) WT injured, (C), LIF Het sham, and (D) LIF Het injured mice. Arrows indicate Olig2+/Ki-67+ cells. (E) Stereological counts of Ki-67+/Olig2+ cells in the SVZ from LIF Het CHI, LIF Het sham, WT sham, and WT CHI mice. n=5 animals per group. ****p≤0.0001 vs. WT injured; WT sham, & LIF Het sham by one-way ANOVA and Tukey’s *post hoc* test. (F-G) Representative images of the WM 2 DPI from (F) LIF Het injured and (G) WT injured mice. (H) Stereological counts of Ki-67+/Olig2+ cells in the WM of LIF Het CHI compared with WT CHI mice. n=5 animals per group. ***p≤0.001 vs. WT injured by one-way ANOVA and Tukey’s *post hoc* test. (I) Quantification of Ki-67+/Olig2+ cells along the rostro-caudal extent of the CHI. n=5 animals per group. ****p≤0.0001 vs. WT injured and **p≤0.01 by one-way ANOVA and Tukey’s *post hoc* test.

As LIF Het injured mice had more Ki-67+/Olig2+ staining than WT injured mice in the SVZ, we also examined the subcortical WM to determine how the regions directly below the impact zone responded. High numbers of Ki-67+/Olig2+ cells were observed in the corpus callosum (CC) residing between the two SVZ’s of the LIF Het injured mice (Fig. 2F), directly below the injury region (Fig. 2G). Stereological quantification of Ki-67+/Olig2+ cells in the LIF Het injured mice and WT injured mice revealed a 2-fold increase of double positive cells in the CC of LIF Het injured mice when compared with WT injured mice (p≤0.001) (Fig. 2I).

To further examine the diffusivity of the injury on proliferating oligodendrocyte progenitors in the SVZ, we examined the breakdown of the extent of Olig2+/Ki-67+ at three different levels: −0.94 mm Bregma, −0.46 mm Bregma, & 0.02 mm Bregma (Fig 2H). Double positive cells were two times more frequent in the more posterior portions of the SVZ of LIF Het injured mice (****p≤0.0001) and the differences between the WT injured and LIF Het injured groups decreased as the distance from the impact site increased (more anterior) at −0.46 mm Bregma (**p≤0.01) and 0.02 mm Bregma (not significant.) These data suggest that oligodendrocyte progenitors re-enter the cell cycle in the LIF Het injured mice post CHI in both the SVZ and WM region.

### SVZ progenitor proliferation is significantly higher in LIF Het mice, especially within the oligodendrocyte lineage

We next used multicolor flow cytometry to examine the specific cell types of the SVZ, with and without injury, across the two genotypes of interest. This flow panel enables the cells of the SVZ to identified as multipotential, bipotential or unipotential progenitors (Buono et al., 2012). P20 mice received a CHI halfway between Bregma and Lambda and were provided drinking water supplemented with EdU, ad libitum, for two days following the injury (Fig, 3A). Two days after CHI, the two SVZs from each animal were microdissected, dissociated and incubated with the antibodies listed in Table 1 and processed for EdU incorporation using Click-IT chemistry. This analysis revealed a 4-fold increase in the proliferating multipotential progenitor-4 cells (MP4) (CD133+/CD15+/NG2+/PDGFRa+/EdU+) of the SVZ in the LIF Het injured mice when compared to sham-operated controls (**p≤0.01) (Fig. 3B) and a 2.5-fold increase in the LIF Het injured mice compared to WT injured mice (*p≤0.05) (Fig. 3B). Additionally, proliferating PDGFRα-FGF2-responsive multipotential cells (PFMP) (CD133-/CD15+/NG2+/CD140+/EdU+) were increased 12.5-fold in the LIF Het injured mice when compared to sham-operated controls (**p≤0.01) and 5-fold when compared to WT injured mice (*p≤0.05) (Fig. 3C). Furthermore, multipotential progenitor-3 (MP3)/glial restricted progenitors-2 (GRP2) cells (CD133-/CD15-/NG2+/CD140-/EdU+) were 2.5 times higher in the LIF Het injured mice when compared to sham-operated controls (**p≤0.01) and 1.6 times higher compared to the WT injured mice (p=0.08) (Fig. 3D). There was no change in EdU incorporation into the NSCs after the injury between genotypes (Tables 2 and 3).

**Figure 3.**
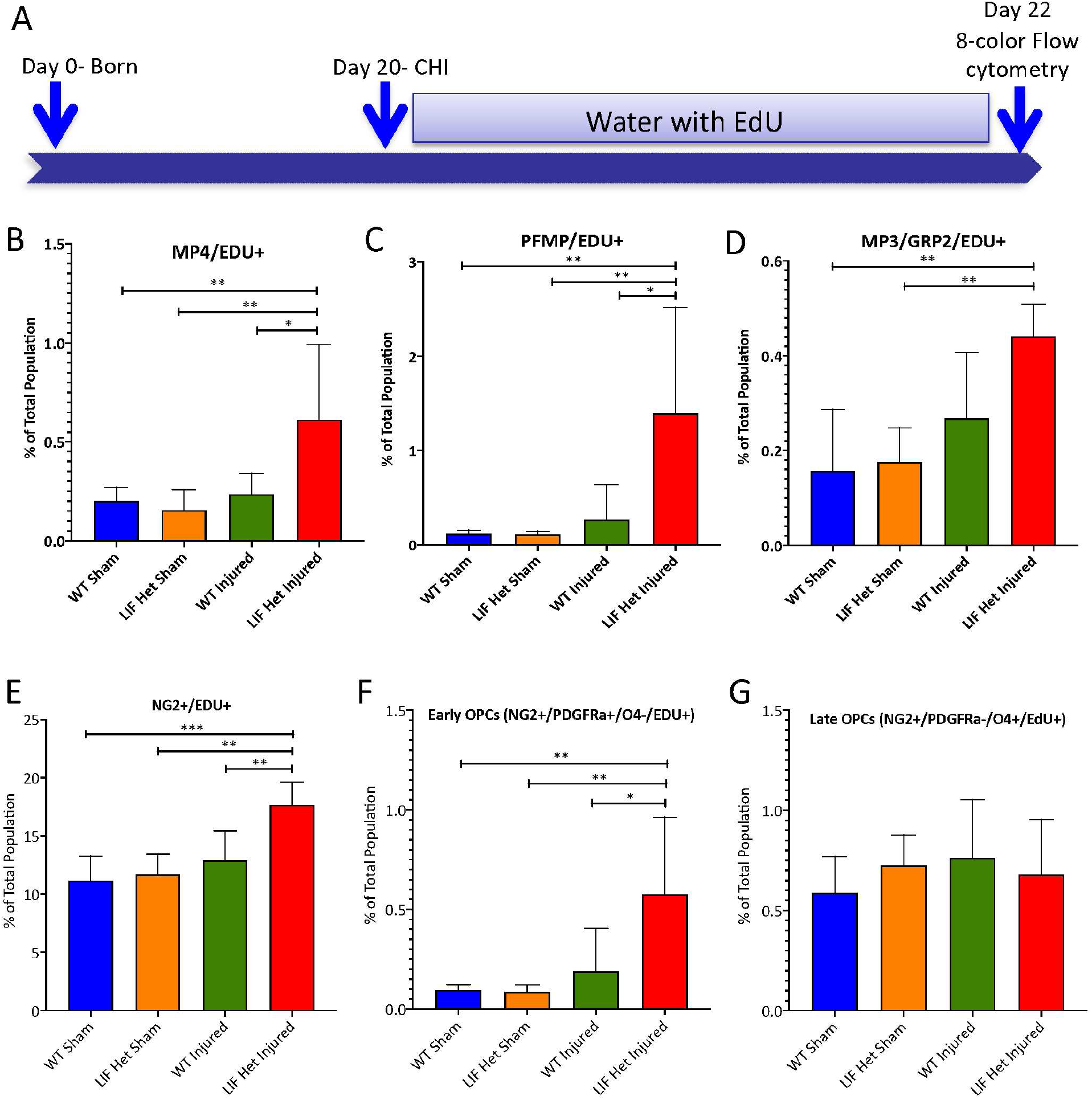
SVZ Progenitor proliferation is significantly higher in LIF Het mice, especially within the oligodendrocyte lineage, as measured by flow cytometry. (A) Schematic depicting experimental design. WT and LIF Het mice were injured at post-natal Day 20 and EdU (1 mg/mL) was provided in sweetened drinking water for 2 days to mark proliferating cells PI. After 2 days, the two SVZs from each animal were microdissected, dissociated, blocked with FcR blocker and stained according to the markers in Table 1. Graphs depict (B) EdU+ MP4 (C) EdU+ PFMP (D) EdU+ MP3/GRP2 cells (E) EdU+ NG2+ cells, (F) EdU+ early OPCs and (G) EdU+ late OPCs. Values represent means ± SDs from at least 5 animals per group. All tests performed by one-way ANOVA and Tukey’s *post hoc* test. * p < 0.05; ** p < 0.01; *** p < 0.001.

**Table 2:** Average percentages of proliferating, SVZ neural stem cells and progenitors at 48 h PI. WT and LIF Het mice were injured at P20. EdU (100 mg/kg) was injected IP at 4 h and 2 h prior to euthanizing to mark proliferating cells post injury. Two days after CHI, the two SVZs from each animal were microdissected, dissociated, blocked with FcR blocker and stained according to the markers in Table 1. Averages are shown as a percent of the total population. Comparisons are noted with the p value between the groups. n= at least 4 animals per group. * p < 0.05 by one-way ANOVA and Tukey’s *post hoc* test.

|  | EdU+ |  |  |  |  |  |  |  |
| --- | --- | --- | --- | --- | --- | --- | --- | --- |
| Group | NSC | MP1 | MP2 | BNAP/GRP1 | MP4 | MP3/GRP2 | PFMP | GRP3 |
| WT injured Avg. | 0.034 | 0.005 | 0.779 | 0.072 | 0.089 | 0.256 | 0.014 | 0.008 |
| WT sham Avg. | 0.033 | 0.004 | 0.649 | 0.810 | 0.094 | 0.178 | 0.013 | 0.005 |
| LIF Het injured Avg. | 0.006 | 0.002 | 1.476 | 0.235 | 0.185 | 0.062 | 0.016 | 0.001 |
| LIF Het sham Avg. | 0.008 | 0.022 | 1.604 | 0.264 | 0.199 | 0.110 | 0.018 | 0.001 |
| Statistical Comparisons | p values |  |  |  |  |  |  |  |
| WT injured vs. WT sham | >0.999 | 0.934 | 0.981 | 0.999 | 0.999 | 0.884 | 0.999 | 0.734 |
| WT injured vs. LIF Het injured | 0.012* | 0.244 | 0.278 | 0.091 | 0.215 | 0.365 | 0.915 | 0.192 |
| WT injured vs. LIF Het sham | 0.021* | 0.441 | 0.161 | 0.039* | 0.133 | 0.595 | 0.671 | 0.262 |
| WT sham vs. LIF Het injured | 0.019* | 0.542 | 0.183 | 0.136 | 0.283 | 0.771 | 0.881 | 0.691 |
| WT sham vs. LIF Het sham | 0.032* | 0.779 | 0.104 | 0.062 | 0.183 | 0.941 | 0.629 | 0.798 |
| LIF Het injured vs. LIF Het sham | 0.993 | 0.979 | 0.989 | 0.975 | 0.993 | 0.981 | 0.968 | 0.998 |

**Table 3:** Average percentage glial progenitors at 48 h PI. WT and LIF Het mice were injured at P20. EdU (100 mg/kg) was injected IP at 4 h and 2 h prior to euthanizing to mark proliferating cells post injury. Two days after CHI, the two SVZs from each animal were microdissected, dissociated, blocked with FcR blocker and stained according to the markers in Table 1. Averages are shown as a percent of the total population. Comparisons are noted with the p value between the groups. n= at least 4 animals per group. * p < 0.05 by one-way ANOVA and Tukey’s *post hoc* test.

|  | **EdU+** |  |  |  |  |  |
| --- | --- | --- | --- | --- | --- | --- |
| **Group** | **O4+** | **O4-/CD133+** | **Glast+** | **Glast+/CD133-** | **Glast+/CD15+** | **Glast+/CD15-** |
| **WT injured Avg.** | 0.835 | 0.264 | 3.579 | 0.553 | 0.908 | 0.284 |
| **WT sham Avg.** | 0.884 | 0.207 | 2.952 | 0.492 | 0.897 | 0.181 |
| **LIF Het injured Avg.** | 3.689 | 0.195 | 5.966 | 1.900 | 4.428 | 0.043 |
| **LIF Het sham Avg.** | 3.615 | 0.230 | 6.149 | 1.988 | 3.680 | 0.071 |
| **Statistical Comparisons** | **p values** |  |  |  |  |  |
| **WT injured vs. WT sham** | >0.999 | 0.478 | 0.928 | 0.999 | >0.999 | 0.841 |
| **WT injured vs. LIF Het injured** | 0.024* | 0.369 | 0.176 | 0.059 | 0.014* | 0.302 |
| **WT injured vs. LIF Het sham** | 0.028* | 0.843 | 0.133 | 0.042* | 0.061 | 0.402 |
| **WT sham vs. LIF Het injured** | 0.033* | 0.992 | 0.078 | 0.057 | 0.018* | 0.749 |
| **WT sham vs. LIF Het sham** | 0.039* | 0.949 | 0.058 | 0.041* | 0.072 | 0.851 |
| **LIF Het injured vs. LIF Het sham** | 0.999 | 0.862 | 0.999 | 0.998 | 0.901 | 0.997 |

Data from Figure 2 suggested that the proliferating cells in LIF Het mice post injury belong to the oligodendrocyte lineage. However, Olig2 is also expressed in a subset of multipotential progenitors within the SVZ (Inamura et al., 2011). Therefore, to establish whether bona fide OPCs are proliferating after the CHI and to determine which stage of the oligodendrocyte lineage is most affected by LIF deficiency and injury, we incorporated specific markers for oligodendrocyte lineage into the flow cytometry panel. By flow cytometry, LIF Het injured mice showed a 50% increase in NG2+/EdU+ cells when compared to WT sham controls (***p≤0.001) and LIF Het sham controls (**p≤0.01) and a 30% increase when compared to WT injured mice (**p≤0.01) (Fig. 3E). However, multipotential progenitors in the SVZ also have been shown to express NG2 (Aguirre et al., 2004). Therefore, we analyzed the percentage of early OPCs (NG2+/PDGFRa+/O4-/EdU+) and found that LIF Het injured mice had a 6-fold increase compared to sham-operated LIF Het controls (**p<0.01) and a 3-fold increase compared to WT injured mice (*p≤0.05) (Fig. 3F). Then, we analyzed the proportion of late OPCs (NG2+/PDGFRa-/O4+/EdU+); however, we did not note differences among the groups (Fig. 3G). These data support the conclusion that a significant proportion of the proliferating cells in the SVZ of LIF Het injured mice are oligodendrocyte progenitors and more specifically, are early OPCs.

### LIF reduces the proliferation of neural progenitors and especially late OPCs

The above studies suggested that LIF was repressing cell proliferation in the SVZ. To establish whether this was a direct effect of LIF on the stem cells and progenitors, we performed an *in vitro* experiment using neurospheres to determine whether LIF would inhibit growth factor stimulated proliferation. To obtain robust data, the cells needed to be synchronized in the G0 stage of the cell cycle prior to providing stimuli. In order to synchronize the cells, they need to be deprived of growth factors for a period of time long enough to arrest their cycling while retaining their ability to reenter the cell cycle.

For these studies secondary neurospheres that had reached ∼200 μm in diameter were deprived of growth factors for 5, 10 or 15 h. The growth factors (EGF & FGF-2) were re-introduced to the cultures along with EdU and after 24 h the cells were analyzed by flow cytometry (Fig. 4A). This experiment revealed that 15 h of growth factor deprivation produced the largest proportion of cells that had been arrested and could re-enter the cell cycle (Fig. 4B).

**Figure 4.**
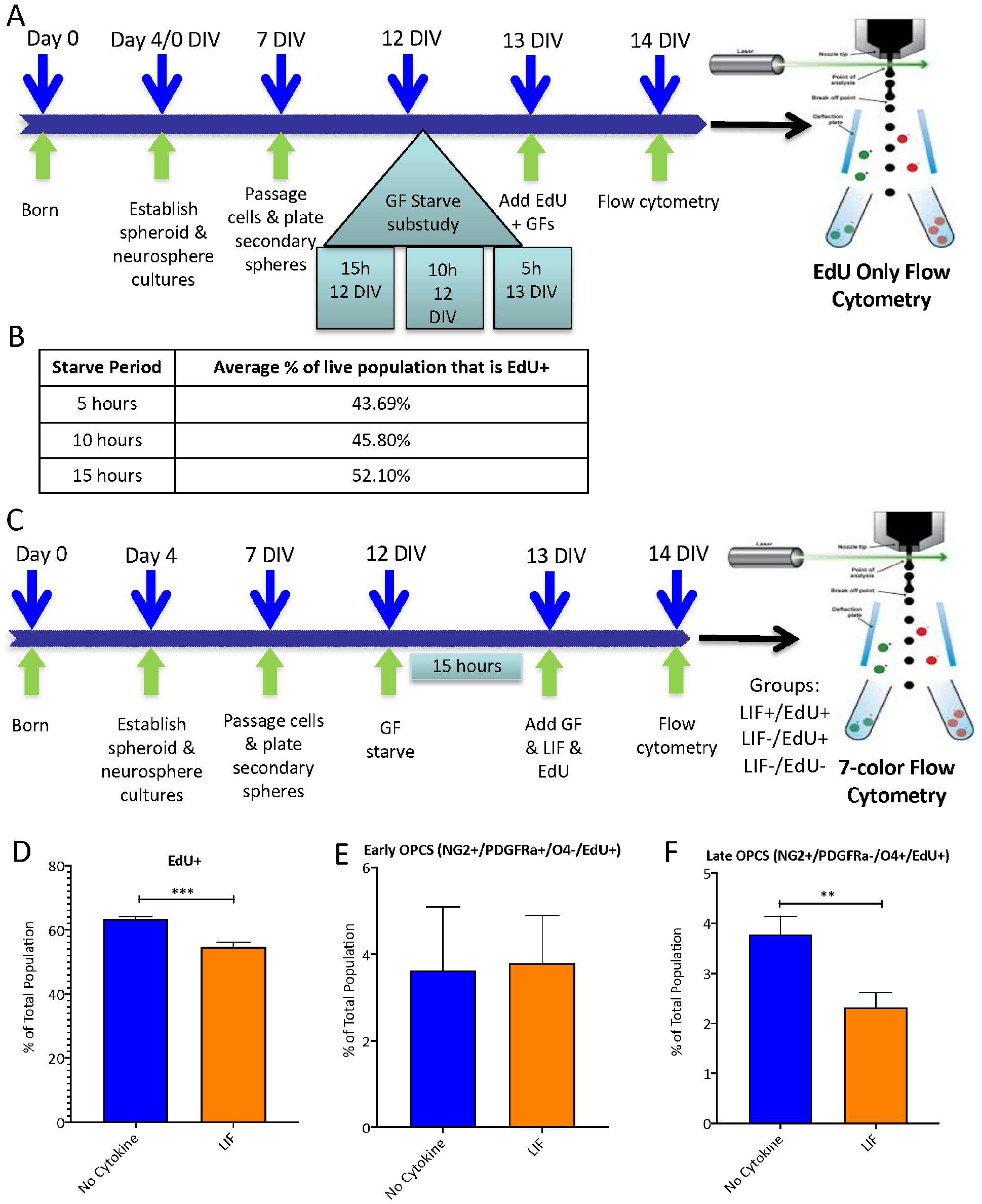
LIF reduces the proliferation of neurospheres, in vitro, and reduces the number of late OPCs as measured by flow cytometry. (A) Schematic of experimental design. Neurospheres were produced from P4 WT mouse SVZs and propagated in growth medium containing 20 ng/mL EGF & 10 ng/mL FGF at 37°C in 5% CO_2_ incubator. After 7 DIV, the primary neurospheres were passaged to generate secondary neurospheres. After 6-7 days, the secondary neurospheres were deprived of growth factors for 15, 10, or 5 h to evaluate cell cycle synchronization. The growth factors were re-introduced to cell cultures and EdU was added. Twenty-four h later, the cells were stained for EdU and assessed using multimarker flow cytometry. (B) Table summarizing the starve periods and percentages of the live population that were EdU+. The 15 h starve period showed the highest percentage of EdU+ cells of the live population when compared to the 5 h & 10 h starve periods. (C) At 12 DIV, all plates were deprived of growth factors for 15 h to enhance cell cycle synchronization. Growth factors (EGF & FGF-2) were re-introduced to cell cultures as well as 5 ng/mL recombinant murine LIF and EdU. After 24 h, samples were processed and stained for flow cytometric analysis. (D) Graph depicting EdU incorporation into all neurosphere cells ± LIF. (E) Graph depicting EdU incorporation into the early oligodendrocyte progenitor population. (F) Graph depicting EdU incorporation into the late OPCs when compared to the control group. Values represent means ± SDs with an n = 3 independent cultures per group. All comparisons were performed using student’s t-test. ** p < 0.01; *** p < 0.001.

Since we’ve seen from previous Ki-67 data (Fig. 1C) that a decrease in LIF increases the number of proliferating cells, we hypothesized that LIF was inhibiting normal proliferation of the cells in the SVZ. To test this hypothesis, we evaluated the effect of LIF on cell proliferation. Secondary neurospheres were deprived of growth factors for 15 h to synchronize the cell cycles whereupon the growth factors ± recombinant murine LIF and EdU were added, followed by multi-color flow cytometry (Fig. 4C). Supporting our hypothesis, there were 15% fewer EdU+ cells when LIF was present (*** p < 0.001) (Fig. 4D). Next, we evaluated the effects of LIF on the proliferation of the early and late OPC populations. LIF had no effect on the early OPC population (Fig. 4E) but reduced the proliferation of the late OPCs by ∼40% when compared to the control group (**p≤0.01) (Fig. 4F, Tables 4 and 5). These data demonstrate that LIF exerts an anti-proliferative effect on the OPCs, consistent with previous data showing that it can push OPCs into a more mature state (Buono et al., 2015; Mayer et al., 1994).

**Table 4:** Average percentages of proliferating NSCs and progenitors after 24 h of LIF treatment or no cytokine treatment. Neural tissue rich in SVZ cells were collected from P4 pups and neurosphere cultures were established. Spheres were propagated for several days, passaged, and grown for an additional 5 days. Spheres were starved of growth factors for 15 h and growth factors were reintroduced along with EdU and 5 ng/mL LIF or no cytokine. After 24 h of treatment, cells were dissociated and stained according to the markers in Table 1 and assessed using flow cytometry. Averages are shown as a percent of the total population. Comparisons are noted with the p value between the groups. n= at least 3 cultures per group. ** p < 0.01 by student’s t test and Tukey’s *post hoc* test.

|  | EdU+ |  |  |  |  |  |  |  |
| --- | --- | --- | --- | --- | --- | --- | --- | --- |
| Group | NSC | MP1 | MP2 | BNAP/GRP1 | MP4 | MP3/GRP2 | PFMP | GRP3 |
| No Cytokine Avg. | 0.004 | 0.070 | 0.020 | 0.301 | 0.237 | 36.746 | 0.321 | 13.592 |
| LIF Avg. | 0.004 | 0.056 | 0.049 | 0.324 | 0.216 | 29.039 | 0.182 | 10.358 |
| <u>Statistical Comparison</u> | <u>p values</u> |  |  |  |  |  |  |  |
| No Cytokine vs. LIF | 0.964 | 0.610 | 0.227 | 0.823 | 0.933 | 0.007** | 0.151 | 0.054 |

**Table 5:**
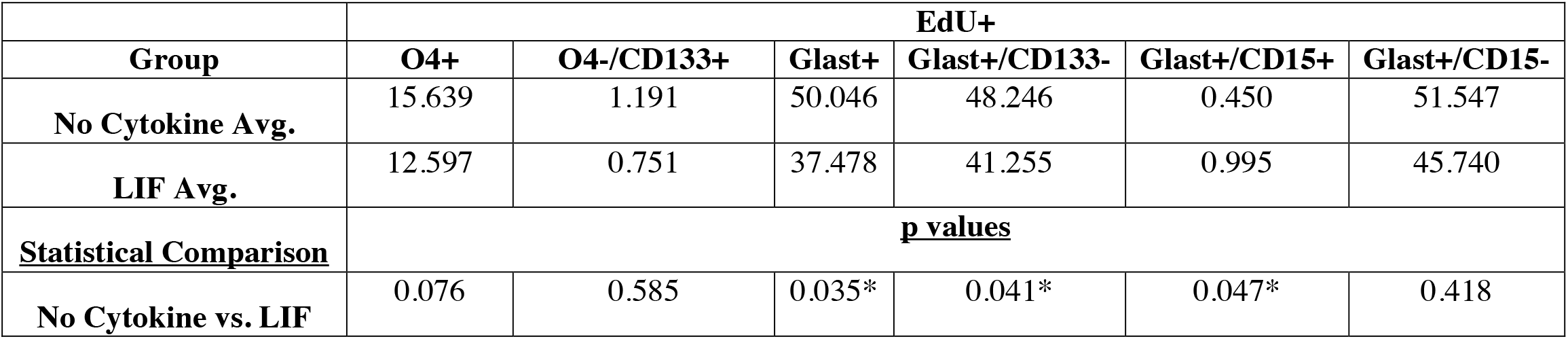
Average percentages of proliferating neural cells and their corresponding P values after 24 h of LIF treatment or no cytokine treatment. Neural tissue rich in SVZ cells were collected from P4 pups and neurosphere cultures were established. Spheres were propagated for several days, passaged, and grown for an additional 5 days. Spheres were starved of growth factors for 15 h and growth factors were reintroduced along with EdU and 5 ng/mL LIF or no cytokine. After 24 h of treatment, cells were dissociated and stained according to the markers in Table1 and assessed using flow cytometry. Averages are shown as a percent of the total population. Comparisons are noted with the p value between the groups. n= at least 3 cultures per group. * p < 0.05 by student’s t test and Tukey’s *post hoc test*.

|  | **EdU+** |  |  |  |  |  |
| --- | --- | --- | --- | --- | --- | --- |
| **Group** | **O4+** | **O4-/CD133+** | **Glast+** | **Glast+/CD133-** | **Glast+/CD15+** | **Glast+/CD15-** |
| **No Cytokine Avg.** | 15.639 | 1.191 | 50.046 | 48.246 | 0.450 | 51.547 |
| **LIF Avg.** | 12.597 | 0.751 | 37.478 | 41.255 | 0.995 | 45.740 |
| **<u>Statistical Comparison</u>** | **<u>p values</u>** |  |  |  |  |  |
| **No Cytokine vs. LIF** | 0.076 | 0.585 | 0.035* | 0.041* | 0.047* | 0.418 |

### LIF activates the mTOR pathway in cultured OPCs

The above studies suggested that LIF could be enhancing the maturation of oligodendrocytes and one of the key pathways associated with oligodendrocyte maturation is the mammalian target of rapamycin (mTOR) pathway. Through both genetic and pharmacological manipulations of this pathway, it has become apparent that mTOR is a key signal transduction pathway for oligodendrocyte differentiation (Ryskalin et al., 2017; Tyler et al., 2009; Wood et al., 2013). Moreover, mTOR regulates oligodendrocyte differentiation during the transition from the late progenitor to the immature oligodendrocyte (Ornelas et al., 2020). Therefore, we hypothesized that LIF might be stimulating the mTOR pathway. As insulin like growth factor-1 (IGF-1) induces oligodendrocyte differentiation, we used IGF-1 as a positive control to measure against LIF for OPC maturation(Wheeler & Fuss, 2016). To test this hypothesis, we prepared highly enriched OPC cultures from newborn mouse forebrain and treated the OPCs with one of four treatment conditions: unstimulated media, 20 ng/mL IGF-1, 5 ng/mL LIF for 20 m, or LIF for 40 m. The samples were collected and analyzed by Western blot for mTOR pathway indicators, pAkt or pS6RP (Fig. 5A). Antibodies to phosphorylated mTOR are not robust, therefore, we did not quantify mTOR phosphorylation itself. Quantification of pAkt (a 60 kDa protein that is upstream of mTOR) revealed that LIF maximally stimulated AKT phosphorylation at 20 m of exposure, resulting in a 2-fold increase when compared to control (Fig. 5B, C). Phosphorylation of AKT had returned to control levels by 40 m of stimulation. Quantification of pS6RP (a 32 kDa protein that is downstream of mTOR) indicated a time dependent increase in phosphorylation with a 3.5-fold increase over the negative control after 20 m of LIF treatment and a 5.8-fold increase after 40 m of LIF exposure (Fig. 5D, E). These data indicate that LIF activates the mTOR pathway and provide mechanistic insight into how LIF can enhance OPC maturation.

**Figure 5.**
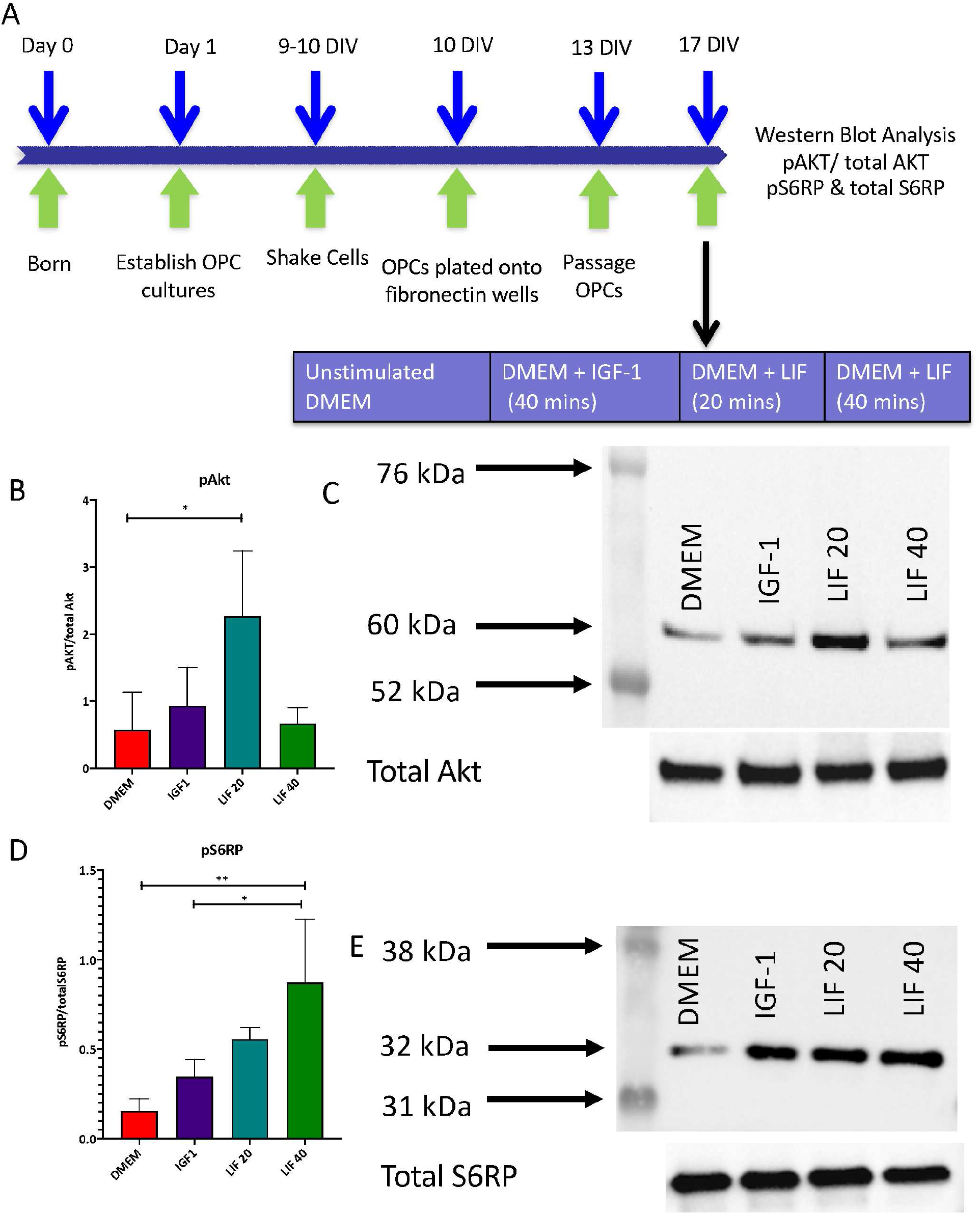
LIF activates the mTOR pathway through pAkt and pS6RP in cultured OPCs. (A) Schematic of experimental design. Mixed glial cultures were prepared from WT P1 mice and then shaken twice after reaching confluence; once to remove microglia and again to remove adherent OPCs. The OPCs were replated onto fibronectin dishes and then passaged 3 days later. OPCs receive one of four treatment conditions: unstimulated media, 20 ng/mL IGF-1, 5 ng/mL rmLIF for 20 m, or LIF for 40 m. The samples were then collected and analyzed by western blot for mTOR pathway indicators, pAkt or pS6RP. (B) Quantification of pAkt from western blot for DMEM (negative control), IGF-1 (positive control), LIF treatment (20 m), & LIF treatment (40 m) for OPCs. (C) Representative image of western blot bands for pAkt at 60 kDa. (D) Quantification of pS6RP from western blot for DMEM (negative control), IGF-1 (positive control), LIF treatment (20 m), & LIF treatment (40 m) for OPCs. (E) Representative image of western blot bands for pAkt at 60 kDa. Graphs represent means ± SD from 3 independent cultures per group. Data were analyzed by one-way ANOVA and Tukey’s *post hoc* test. * p < 0.05; ** p < 0.01.

## Discussion

In the experiments described in this paper, we used several complementary approaches to determine the changes in cell proliferation following a mild CHI, as well as how LIF deficiency affects which cells are proliferating. We produced a mild CHI to the neocortex of P20 mice and performed immunofluorescence and flow cytometric analyses to identify and quantify the distinct SVZ neural progenitors. Our studies demonstrated that a variety of neural progenitors are stimulated to proliferate including multipotential, bipotential and uni-potential progenitors. Given the vulnerability of the WM in concussive injuries, we focused on evaluating the progenitors of the oligodendrocyte lineage and show that reduced levels of LIF during recovery from brain injury increased the numbers of newly generated oligodendrocyte progenitors. Early OPCs proliferated 6-fold more, as revealed by flow cytometry and there were 3-fold more proliferating Olig2 expressing cells in the SVZ. Our studies using neurospheres demonstrated that LIF reduces the number of total EdU+ cells, consistent with its role as a niche component that helps to maintain the stem cells and progenitors. LIF also decreased the number of proliferating OPCs, supporting a model where the increase in LIF after injury is suppressing the expansion of OPCs. Applying LIF to highly enriched OPC cultures revealed that LIF activates the AKT/mTOR pathway, consistent with the important role for this pathway in OPC maturation. Altogether, these studies demonstrate that LIF exerts anti-proliferative effects on the neural progenitor cell population, where it can either keep progenitors in a primitive, more quiescent state or push later stage progenitors to terminally differentiate.

Approximately half of all TBIs cause changes in WM integrity (Mac Donald et al., 2011). Within the injured WM, there may be demyelination of intact axons that have the potential to be remyelinated to restore function (Mierzwa et al., 2015). Therefore, a goal of this study was to determine whether the persistent deficits in myelination in the LIF Het mice might be due to a depressed regenerative response from the stem cells and progenitors of the SVZ. In our previous study of CHI, LIF Het mice had persistently reduced myelination after injury. The injured LIF Het mice also presented with more severe motor and sensory deficits, compared with WT injured mice. The prolonged accumulation of neurological impairment was accompanied by an increase in cell death, a decrease in the total number of Olig2+ cells in the WM, axonal degeneration, hypomyelination and a diminished callosal compound action potential, especially of the myelinated axons (Goodus et al., 2016). Since this injury took place prior to the completion of myelination in the juvenile mouse, the reduced myelination in the injured LIF Het mice, suggested that there might be a reduced response by progenitors in the SVZ cells in LIF Het mice that contributed to the decreased recovery of the WM following CHI. Therefore, it was surprising when the LIF Het injured mice demonstrated increased OPC proliferation. Goodus et al., 2016 observed a 50% decrease in myelin basic protein (MBP) staining in the CC (WM) at 2, 7, and 14 days after CHI in LIF Het mice when compared to WT mice. However, a study by Gresle et al., 2012, reported that there was greater axonal injury in LIF deficient mice than WT mice in an EAE model that could be attributed to direct effects on the axons rather than beneficial actions through the oligodendrocytes (Gresle et al., 2012). Furthermore, in a study of CHI in adult mice, Marion et al., 2018 found that OPCs proliferated after the injury, but that they failed to differentiate into mature myelinating oligodendrocytes, leading them to conclude that the failure of remyelination is not due to the absence of pre-myelinating OPCs (Marion et al., 2018). Our studies demonstrating increased OPC proliferation in LIF Het mice at 2 days post CHI suggest that more severe axonal damage in the injured LIF Het mice likely reduced the numbers of axons available for myelination and, therefore, even with increased numbers of OPCs, there was not enough substrate for the OPCs to myelinate. These results provide important insight into the mechanism by which LIF influences injury repair.

Taking into account the drastic increase in proliferation post injury of LIF Het mice SVZ and, more specifically, the increase in oligodendrocyte progenitors in response to injury, it seems that LIF is critical in regulating SVZ progenitor cell proliferation after pediatric TBI. One concern that we had was whether the observed changes in SVZ cell proliferation were a result of LIF acting directly on the progenitors, or whether it was acting indirectly, as the injured cells or recruited macrophages (that have LIF receptors) could have produced diffusible factors that affected the progenitors in the SVZ and subcortical WM. For example, in a model of spinal cord injury, daily injections of LIF to female C57 black mice, inhibited oligodendrocyte death after injury, as measured by a reduction in cleaved caspase-3+ CC1+ cells (Kerr & Patterson, 2005). However, with the LIF injections there were more reactive microglia and increased levels of IGF-1. The authors concluded that LIF was not exerting a direct action on the oligodendrocytes, but rather that LIF was increasing the production and release of IGF-1 from the macrophages, which was promoting the survival of the oligodendrocytes (Kerr & Patterson, 2005). By contrast, Lin et al., 2020, found that IGF-1 levels were higher in injured LIF Het mice after a neonatal hypoxia-ischemic brain injury, which is inconsistent with the proposed indirect actions of LIF through IGF-1 (Lin et al., 2020). Furthermore, arguing against an indirect mechanism, we used a PCR array to screen mRNAs for 85 growth factors and cytokines and did not find any that were differentially expressed by the injured LIF Het neocortex vs. the WT injured cortex (data not shown).

Other studies have demonstrated that brain injuries are worse when there is a deficiency in LIF. Butzkueven et al., 2006 showed that administering neutralizing antibodies to LIF worsened the severity of demyelination and doubled oligodendrocyte loss using an EAE model. Blocking LIF impaired the clinical outcome (as measured by myelin damage) but did not reduce oligodendrocyte progenitor numbers. This suggests that LIF affects mature oligodendrocytes but does not directly affect OPCs (Butzkueven et al., 2006). Similarly, LIF knock out mice experience more demyelination than WT mice, possibly due to the loss of mature oligodendrocytes or failure of progenitors to respond to oligodendrocyte loss (Marriott et al., 2008). Our results show that when LIF is reduced, the OPCs may remain in an immature, proliferative state, whereas applying LIF to OPC cultures pushes them into a more mature state. Goodus et al. 2016 also showed that LIF Het mice had reduced levels of astrogliosis and microgliosis at 2 days post CHI in the CC (Goodus et al., 2016). Thus, one might have predicted that reduced levels of LIF would result in fewer oligodendrocytes, but this was clearly not the case. Altogether these data lead to the conclusion that LIF has complex effects on the oligodendrocyte lineage where it protects mature oligodendrocytes after injury and can also push OPCs into a more mature state to promote the repopulation of the mature oligodendrocyte niche.

Administering LIF promotes oligodendrocyte survival in several injury models. Mayer et al., 1994 applied LIF to rat optic nerve cells and found that LIF increased the generation and maturation of OPCs (Mayer et al., 1994). Similarly, Gresle et al., 2012 found that the addition of LIF (100ng/mL) to oligodendrocyte cultures increased survival after 2 and 3 days of administration when compared to basal media (Gresle et al., 2012). After intranasal administration, Lin et al., 2020 observed increased CC thickness and MBP staining in a model of perinatal hypoxia-ischemia. In that study, they evaluated BrdU incorporation into Olig2+ cells after intranasal LIF treatment and failed to show any differences between vehicle and LIF treated mice (Lin et al., 2020), leading them to conclude that intranasally administered LIF was exerting a strong survival promoting effect on the existing oligodendrocytes.

In a separate study using NG2 targeted LIF loaded nanoparticles to deliver LIF to oligodendrocyte progenitors in a model of focal CNS demyelination, Rittchen et al., 2015 demonstrated that LIF increased Stat-3 phosphorylation in the OPCs and that it promoted their differentiation into mature oligodendrocytes, resulting in increased numbers of myelinated axons and increased myelin thickness (Rittchen et al., 2015) leading them to conclude that LIF promoted oligodendrocyte progenitor cell maturation through Stat-3. However, here we demonstrate that LIF increases AKT and s6RP phosphorylation when applied to OPC cultures, consistent with the view that other pathways besides the Jak-Stat-3 pathway are important for stimulating OPC maturation. As LIF depresses the regenerative response of the progenitors in the SVZ, the data presented here indicate that if LIF is to be used as a therapeutic to promote WM repair, that it be delivered during subacutely after injury.

## Acknowledgements

Research reported in this publication was supported by the New Jersey Commission on Brain Injury Research under award numbers [CBIR19Felo14] to MJF and [CBIR13IRG0170] to SWL.

## Figure Legends

**Supplemental Fig 1.**
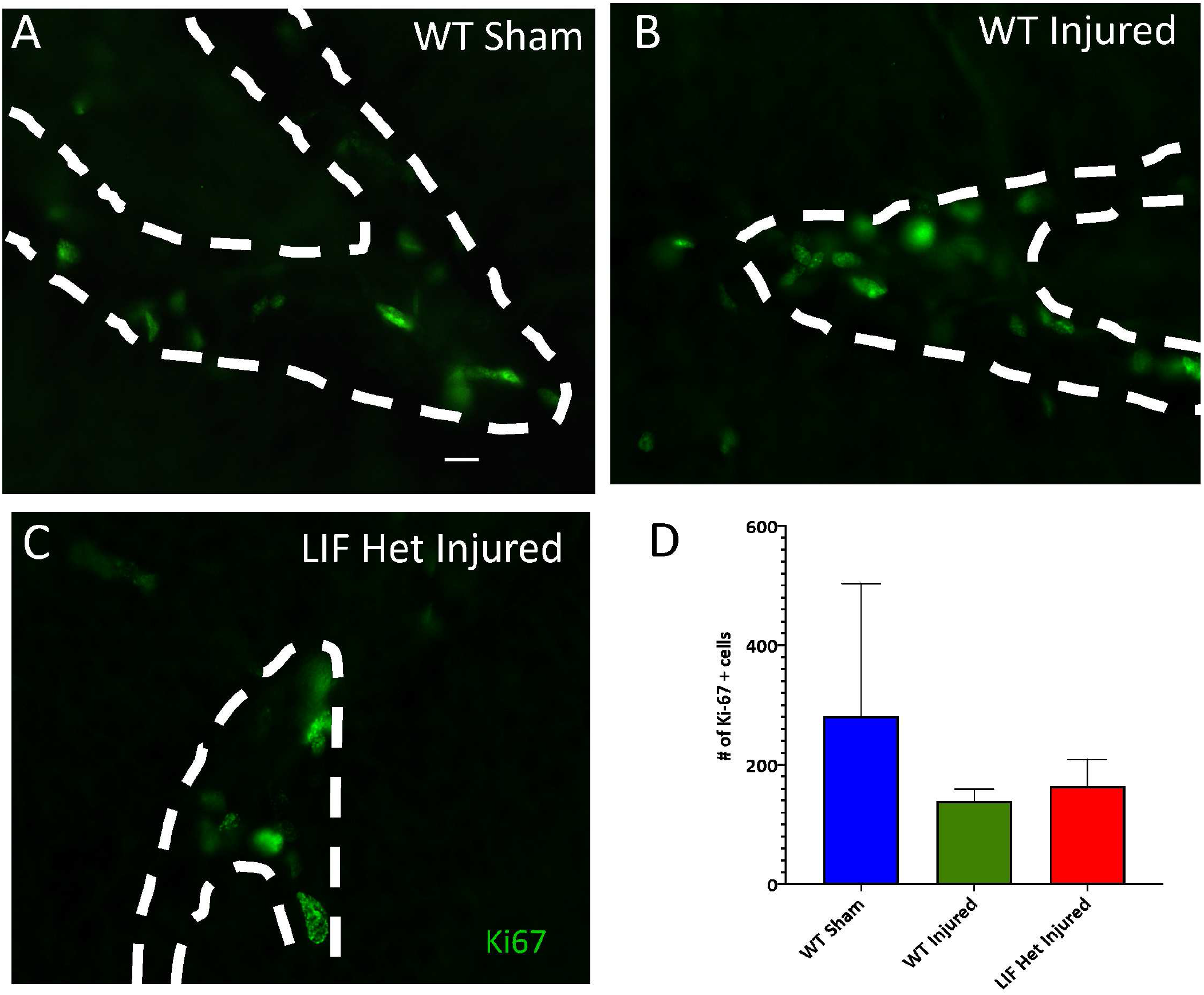
SGZ cell proliferation in LIF Het mice does not differ significantly from WT mice after CHI. WT and LIF Het mice were injured at P20 and analyzed at 2 DPI. Coronal 40 μm sections containing the SGZ were stained for the cell proliferation marker, Ki-67. (A-C) Representative images of the SGZ below the center of the impact zone at 2 DPI from (A) WT sham, (B) WT injured, and (C) LIF Het injured mice. (D) Stereological counts of Ki-67+ cells in the SGZ from WT sham, WT Injured, and LIF Het CHI mice. Data represent the means ± SD from n=3 animals per group. Data were analyzed by one-way ANOVA and Tukey’s post hoc test. Magnification bar = 10 µm.

**Supplemental Figure 2.**
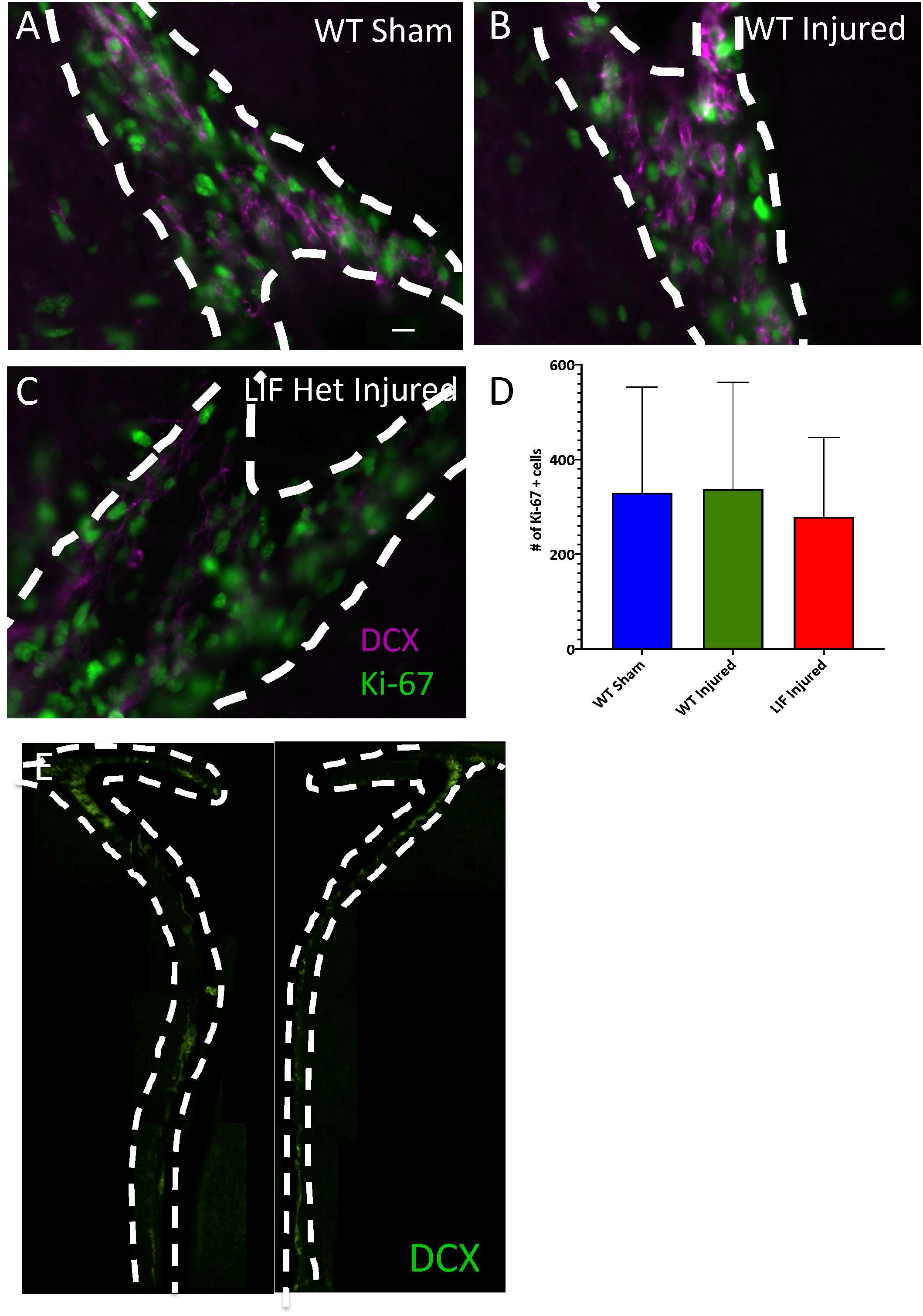
Neuronal precursor population is unchanged at 48h PI. WT and LIF Het mice were injured at post-natal day 20 and analyzed 2 days PI. Coronal 40 μm sections containing the SVZ were stained for Ki-67 and (the neuronal precursor marker) doublecortin (DCX). (A-C) Representative images of the dorsomedial lateral SVZ below the center of the impact zone 2 DPI from (A) WT sham, (B) WT injured, and (C) LIF Het injured mice. (D) Stereological counts of Ki-67+/DCX+ cells in the SVZ from WT sham, WT Injured, and LIF Het CHI mice. Values represent means ± SDs. (E) Whole SVZ images of DCX staining. n=3 animals per group. Analyzed by one-way ANOVA and Tukey’s *post hoc* test.

## Graphical Abstract: LIF Haplodeficiency Alters Forebrain Development and Proliferative Responses to CHI

Injury increases overall proliferation and increases the number of multipotential progenitors in the SVZ at 48 h PI that is more dramatic in LIF Het mice. Oligodendrocyte lineage cells are most sensitive to decreased levels in LIF and more specifically, early OPCs proliferation increases in LIF Het injured mice. Addition of LIF decreases overall proliferation and decreases the number of proliferating late OPCs, suggesting that LIF pushes the cells into a more mature state.

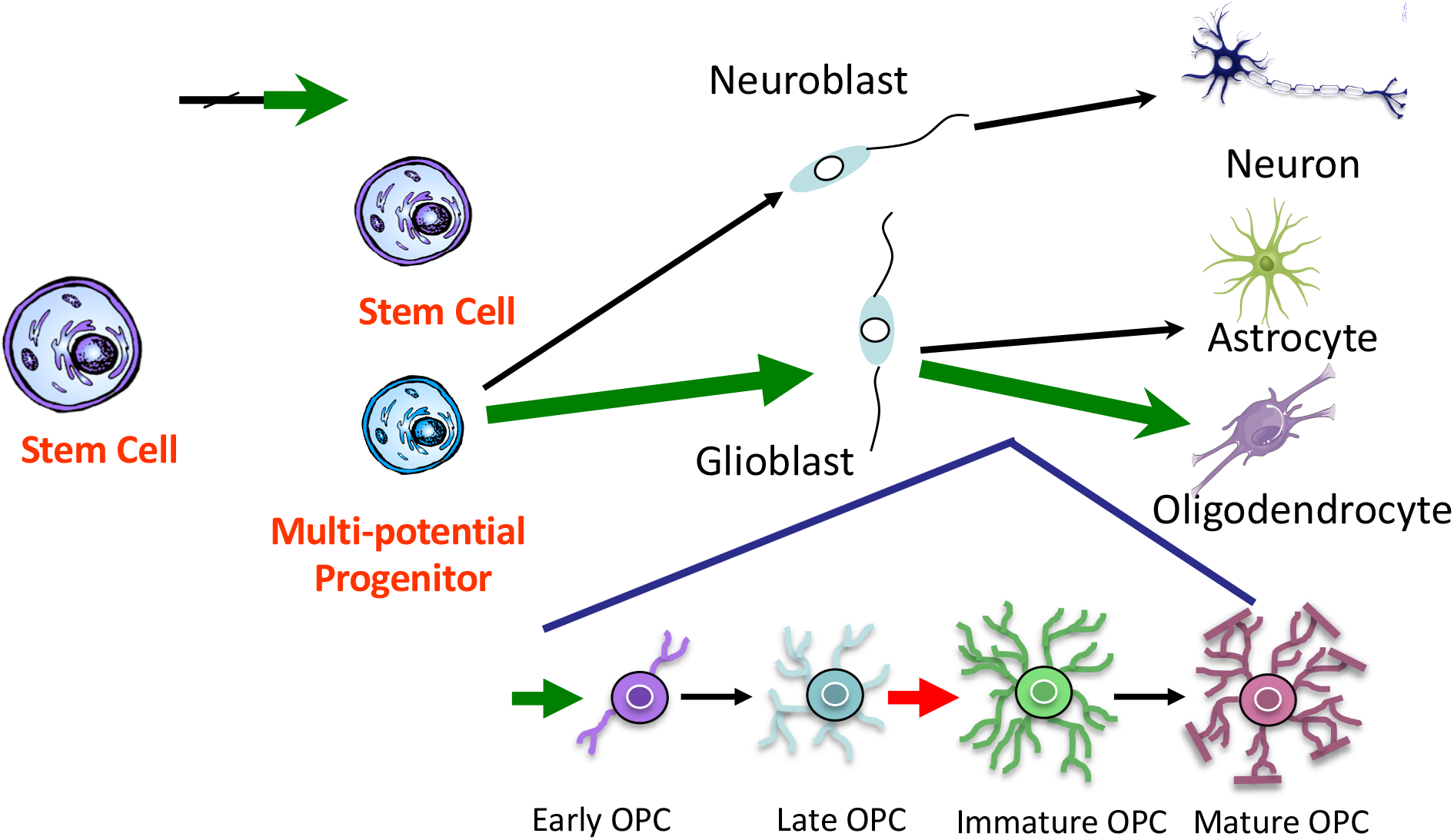

## References Cited

Aguirre, A. A., Chittajallu, R., Belachew, S., & Gallo, V. (2004). NG2-expressing cells in the subventricular zone are type C-like cells and contribute to interneuron generation in the postnatal hippocampus. Journal of Cell Biology, 165(4), 575–589.

Bauer, S., & Patterson, P. H. (2006). Leukemia inhibitory factor promotes neural stem cell self-renewal in the adult brain. J Neurosci, 26(46), 12089–12099.

Buono, K. D., Goodus, M. T., Guardia Clausi, M., Jiang, Y., Loporchio, D., & Levison, S. W. (2015). Mechanisms of mouse neural precursor expansion after neonatal hypoxia-ischemia. J Neurosci, 35(23), 8855–8865. doi:10.1523/JNEUROSCI.2868-12.2015

Buono, K. D., Vadlamuri, D., Gan, Q., & Levison, S. W. (2012). Leukemia inhibitory factor is essential for subventricular zone neural stem cell and progenitor homeostasis as revealed by a novel flow cytometric analysis. Dev Neurosci, 34(5), 449–462. doi:10.1159/000345155

Butzkueven, H., Emery, B., Cipriani, T., Marriott, M. P., & Kilpatrick, T. J. (2006). Endogenous leukemia inhibitory factor production limits autoimmune demyelination and oligodendrocyte loss. Glia, 53(7), 696–703.

Centers for Disease Control and Prevention (CDC), N. C. f. I. P. a. C. (2019). Severe TBI. https://www.cdc.gov/traumaticbraininjury/severe.html

Covey, M. V., Jiang, Y., Alli, V. V., Yang, Z., & Levison, S. W. (2010). Defining the critical period for neocortical neurogenesis after pediatric brain injury. Dev Neurosci, 32(5-6), 488–498. doi:10.1159/000321607

Deverman, B. E., & Patterson, P. H. (2012). Exogenous leukemia inhibitory factor stimulates oligodendrocyte progenitor cell proliferation and enhances hippocampal remyelination. J Neurosci, 32(6), 2100–2109. doi:10.1523/JNEUROSCI.3803-11.2012

Faiz, M., Acarin, L., Castellano, B., & Gonzalez, B. (2005). Proliferation dynamics of germinative zone cells in the intact and excitotoxically lesioned postnatal rat brain. BMC Neurosci, 6, 26. doi:10.1186/1471-2202-6-26

Felling, R. J., Covey, M. V., Wolujewicz, P., Batish, M., & Levison, S. W. (2016). Astrocyte-produced leukemia inhibitory factor expands the neural stem/progenitor pool following perinatal hypoxia-ischemia. J Neurosci Res, 94(12), 1531–1545. doi:10.1002/jnr.23929

Goodus, M. T., Guzman, A. M., Calderon, F., Jiang, Y., & Levison, S. W. (2015). Neural stem cells in the immature, but not the mature, subventricular zone respond robustly to traumatic brain injury. Dev Neurosci, 37(1), 29–42. doi:10.1159/000367784

Goodus, M. T., Kerr, N. A., Talwar, R., Buziashvili, D., Fragale, J. E., Pang, K. C., & Levison, S. W. (2016). Leukemia Inhibitory Factor Haplodeficiency Desynchronizes Glial Reactivity and Exacerbates Damage and Functional Deficits after a Concussive Brain Injury. J Neurotrauma, 33(16), 1522–1534. doi:10.1089/neu.2015.4234

Gresle, M. M., Alexandrou, E., Wu, Q., Egan, G., Jokubaitis, V., Ayers, M.,… Butzkueven, H. (2012). Leukemia inhibitory factor protects axons in experimental autoimmune encephalomyelitis via an oligodendrocyte-independent mechanism. PLoS One, 7(10), e47379. doi:10.1371/journal.pone.0047379

Inamura, N., Ono, K., Takebayashi, H., Zalc, B., & Ikenaka, K. (2011). Olig2 lineage cells generate GABAergic neurons in the prethalamic nuclei, including the zona incerta, ventral lateral geniculate nucleus and reticular thalamic nucleus. Dev Neurosci, 33(2), 118–129. doi:10.1159/000328974

Jinnou, H., Sawada, M., Kawase, K., Kaneko, N., Herranz-Perez, V., Miyamoto, T.,… Sawamoto, K. (2018). Radial Glial Fibers Promote Neuronal Migration and Functional Recovery after Neonatal Brain Injury. Cell Stem Cell, 22(1), 128–137 e129. doi:10.1016/j.stem.2017.11.005

Kernie, S. G., & Parent, J. M. (2010). Forebrain neurogenesis after focal Ischemic and traumatic brain injury. Neurobiology of Disease, 37(2), 267–274. doi:10.1016/j.nbd.2009.11.002

Kerr, B. J., & Patterson, P. H. (2005). Leukemia inhibitory factor promotes oligodendrocyte survival after spinal cord injury. Glia, 51(1), 73–79.

Lin, J., Niimi, Y., Clausi, M. G., Kanal, H. D., & Levison, S. W. (2020). Neuroregenerative and protective functions of Leukemia Inhibitory Factor in perinatal hypoxic-ischemic brain injury. Exp Neurol, 113324. doi:10.1016/j.expneurol.2020.113324

Lois, C., & Alvarez-Buylla, A. (1993). Proliferating subventricular zone cells in the adult mammalian forebrain can differentiate into neurons and glia. Proc.Natl.Acad.Sci.USA., 90, 2074–2077.

Mac Donald, C. L., Johnson, A. M., Cooper, D., Nelson, E. C., Werner, N. J., Shimony, J. S.,… Brody, D. L. (2011). Detection of blast-related traumatic brain injury in US military personnel. N Engl J Med, 364(22), 2091–2100. doi:10.1056/NEJMoa1008069

Marion, C. M., Radomski, K. L., Cramer, N. P., Galdzicki, Z., & Armstrong, R. C. (2018). Experimental Traumatic Brain Injury Identifies Distinct Early and Late Phase Axonal Conduction Deficits of White Matter Pathophysiology, and Reveals Intervening Recovery. J Neurosci, 38(41), 8723–8736. doi:10.1523/JNEUROSCI.0819-18.2018

Marriott, M. P., Emery, B., Cate, H. S., Binder, M. D., Kemper, D., Wu, Q.,… Kilpatrick, T. J. (2008). Leukemia inhibitory factor signaling modulates both central nervous system demyelination and myelin repair. Glia, 56(6), 686–698. doi:10.1002/glia.20646

Mayer, M., Bhakoo, K., & Noble, M. (1994). Ciliary neurotrophic factor and leukemia inhibitory factor promote the generation, maturation and survival of oligodendrocytes in vitro. Development, 120(1), 143–153.

Mierzwa, A. J., Marion, C. M., Sullivan, G. M., McDaniel, D. P., & Armstrong, R. C. (2015). Components of myelin damage and repair in the progression of white matter pathology after mild traumatic brain injury. J Neuropathol Exp Neurol, 74(3), 218–232. doi:10.1097/NEN.0000000000000165

Ong, J., Plane, J. M., Parent, J. M., & Silverstein, F. S. (2005). Hypoxic-ischemic injury stimulates subventricular zone proliferation and neurogenesis in the neonatal rat. Pediatr Res, 58(3), 600–606. doi:10.1203/01.PDR.0000179381.86809.02

Ornelas, I. M., Khandker, L., Wahl, S. E., Hashimoto, H., Macklin, W. B., & Wood, T. L. (2020). The mechanistic target of rapamycin pathway downregulates bone morphogenetic protein signaling to promote oligodendrocyte differentiation. Glia, 68(6), 1274–1290. doi:10.1002/glia.23776

Parent, J. M., Vexler, Z. S., Gong, C., Derugin, N., & Ferriero, D. M. (2002). Rat forebrain neurogenesis and striatal neuron replacement after focal stroke. Ann Neurol, 52(6), 802–813.

Plane, J. M., Liu, R., Wang, T. W., Silverstein, F. S., & Parent, J. M. (2004). Neonatal hypoxic-ischemic injury increases forebrain subventricular zone neurogenesis in the mouse. Neurobiology of Disease, 16(3), 585–595. doi:10.1016/j.nbd.2004.04.003

Ramaswamy, S., Goings, G. E., Soderstrom, K. E., Szele, F. G., & Kozlowski, D. A. (2005). Cellular proliferation and migration following a controlled cortical impact in the mouse. Brain Research, 1053(1-2), 38–53. doi:10.1016/j.brainres.2005.06.042

Rittchen, S., Boyd, A., Burns, A., Park, J., Fahmy, T. M., Metcalfe, S., & Williams, A. (2015). Myelin repair in vivo is increased by targeting oligodendrocyte precursor cells with nanoparticles encapsulating leukaemia inhibitory factor (LIF). Biomaterials, 56, 78–85. doi:10.1016/j.biomaterials.2015.03.044

Rowitch, D. H., Lu, Q. R., Kessaris, N., & Richardson, W. D. (2002). An ‘oligarchy’ rules neural development. Trends in Neuroscience, 25(8), 417–422.

Ryskalin, L., Lazzeri, G., Flaibani, M., Biagioni, F., Gambardella, S., Frati, A., & Fornai, F. (2017). mTOR-Dependent Cell Proliferation in the Brain. Biomed Res Int, 2017, 7082696. doi:10.1155/2017/7082696

Semple, B. D., Blomgren, K., Gimlin, K., Ferriero, D. M., & Noble-Haeusslein, L. J. (2013). Brain development in rodents and humans: Identifying benchmarks of maturation and vulnerability to injury across species. Prog Neurobiol, 106-107, 1–16. doi:10.1016/j.pneurobio.2013.04.001

Soilu-Hanninen, M., Broberg, E., Roytta, M., Mattila, P., Rinne, J., & Hukkanen, V. (2010). Expression of LIF and LIF receptor beta in Alzheimer’s and Parkinson’s diseases. Acta Neurol Scand, 121(1), 44–50. doi:10.1111/j.1600-0404.2009.01179.x

Stewart, C. L., Kaspar, P., Brunet, L. J., Bhatt, H., Gadi, I., Kontgen, F., & Abbondanzo, S. J. (1992). Blastocyst implantation depends on maternal expression of leukaemia inhibitory factor. Nature, 359(6390), 76–79. doi:10.1038/359076a0

Susarla, B. T., Villapol, S., Yi, J. H., Geller, H. M., & Symes, A. J. (2014). Temporal patterns of cortical proliferation of glial cell populations after traumatic brain injury in mice. ASN NEURO, 6(3), 159–170. doi:10.1042/AN20130034

Tyler, W. A., Gangoli, N., Gokina, P., Kim, H. A., Covey, M., Levison, S. W., & Wood, T. L. (2009). Activation of the mammalian target of rapamycin (mTOR) is essential for oligodendrocyte differentiation. J Neurosci, 29(19), 6367–6378. doi:29/19/6367 [pii] 10.1523/JNEUROSCI.0234-09.2009

Vanderlocht, J., Hellings, N., Hendriks, J. J., Vandenabeele, F., Moreels, M., Buntinx, M.,… Stinissen, P. (2006). Leukemia inhibitory factor is produced by myelin-reactive T cells from multiple sclerosis patients and protects against tumor necrosis factor-alpha-induced oligodendrocyte apoptosis. J Neurosci Res, 83(5), 763–774.

Wheeler, N. A., & Fuss, B. (2016). Extracellular cues influencing oligodendrocyte differentiation and (re)myelination. Exp Neurol, 283(Pt B), 512–530. doi:10.1016/j.expneurol.2016.03.019

Wood, T. L., Bercury, K. K., Cifelli, S. E., Mursch, L. E., Min, J., Dai, J., & Macklin, W. B. (2013). mTOR: a link from the extracellular milieu to transcriptional regulation of oligodendrocyte development. ASN NEURO, 5(1), e00108. doi:10.1042/AN20120092

